# *Gemin4* is an essential gene in mice, and its overexpression in human cells causes relocalization of the SMN complex to the nucleoplasm

**DOI:** 10.1101/242529

**Authors:** Ingo D. Meier, Michael P. Walker, A. Gregory Matera

## Abstract

Gemin4 is a member of the Survival Motor Neuron (SMN) protein complex, which is responsible for the assembly and maturation of Sm-class small nuclear ribonucleoproteins (snRNPs). In metazoa, Sm snRNPs are assembled in the cytoplasm and subsequently imported into the nucleus. We previously showed that the SMN complex is required for snRNP import *in vitro*, although it remains unclear which specific components direct this process. Here, we report that Gemin4 overexpression drives SMN and the other Gemin proteins from the cytoplasm into the nucleus. Moreover, it disrupts the subnuclear localization of the Cajal body marker protein, coilin, in a dose-dependent manner. We identified three putative nuclear localization signal (NLS) motifs within Gemin4, one of which is necessary and sufficient to direct nuclear import. Overexpression of Gemin4 constructs lacking this NLS sequestered Gemin3 and, to a lesser extent Gemin2, in the cytoplasm but had little effect on the nuclear accumulation of SMN. We also investigated the effects of Gemin4 depletion in the laboratory mouse, *mus musculus. Gemin4* null mice die early in embryonic development, demonstrating that Gemin4 is an essential mammalian protein. When crossed onto a severe SMA mutant background, heterozygous loss of *Gemin4* failed to modify the early postnatal mortality phenotype of SMA type I (*Smn^-/-^;SMN2^+/+^*) mice. We conclude that Gemin4 plays an essential role in mammalian snRNP biogenesis, and may facilitate import of the SMN complex (or subunits thereof) into the nucleus.

## Introduction

Pre-mRNA splicing is a central feature of the eukaryotic gene expression programme. The removal of intronic sequences from pre-mRNAs is catalyzed by a macromolecular machine called the spliceosome. Key components of spliceosomes include the small nuclear ribonucleoproteins (snRNPs). Each of these snRNPs contains a common set of seven RNA binding factors, called Sm proteins, that forms a heptameric ring around the snRNA, known as the Sm core. Biogenesis of the Sm core is carried out by another macromolecular assemblage called the Survival Motor Neuron (SMN) complex, consisting of at least nine proteins (Gemins 2–8, unrip and SMN), reviewed in (Battle *et al*. 2006a; Matera *et al*. 2007; Matera and Wang 2014).

Following RNA polymerase II-mediated transcription in the nucleus, pre-snRNAs are exported to the cytoplasm for assembly into stable RNP particles (Jarmolowski *et al*. 1994; Ohno *et al*. 2000). The SMN complex is thought to bind both the Sm proteins (B, D1, D2, D3, E, F, G) and the uridine-rich snRNAs (U1, U2, U11, U12, etc.), bringing them together and forming the Sm core RNP (Chari *et al*. 2008; Pellizzoni *et al*. 2002). Following 3’ processing and hypermethylation of the snRNA 5’ cap, the Sm core RNP is transported from the cytoplasm back into the nucleus. This process requires one of two known nuclear localization signals (NLSs), the 5’ trimethylguanosine (TMG) cap and the Sm core (Fischer *et al*. 1993; Marshallsay and Luhrmann 1994). The adaptor that recognizes the cap is a protein called Snurportin (Huber *et al*. 1998), whereas the Sm core adaptor is recognized by factor(s) within the SMN complex itself (Narayanan *et al*. 2004). Both adaptors use a common import receptor protein, Importin-β (Huber *et al*. 2002; Palacios *et al*. 1997).

The cytoplasmic SMN complex is thought to chaperone RNP biogenesis by conferring stringent specificity toward snRNAs and preventing illicit Sm core assembly on nontarget RNAs (Chari *et al*. 2008; Pellizzoni *et al*. 2002). The function of the nuclear SMN complex is completely unclear, and Gemin4 is the only member of the complex that contains a classical NLS motif (see below). A number of subcomplexes containing SMN and/or various Gemins have also been described (Battle *et al*. 2007; Hao *et al*. 2007; Yong *et al*. 2010), the functions of which are also unknown. Biochemical and cell biological analyses of the SMN complex have begun to elucidate functions of some of its individual components. SMN and Gemin2 form the core of the complex and are thought to be the primordial proteins in the evolution of the complex (Kroiss *et al*. 2008). Gemin5 is thought to deliver snRNAs to the SMN complex by recognizing specific RNA structural and sequence features including the Sm protein binding site (Battle *et al*. 2006b; Lau *et al*. 2009; Yong *et al*. 2010) and the 7-methylguanosine cap (Bradrick and Gromeier 2009). Gemins 6 and 7 are hypothesized to act as scaffolding intermediates during assembly of the Sm core (Battle *et al*. 2006a; Ma *et al*. 2005). Gemin8 interacts directly with SMN and forms a bridge for the Gemin6/7 heterodimer complex with unrip, bringing this trimeric module into the rest of the SMN complex (Carissimi *et al*. 2005; Carissimi *et al*. 2006a; Carissimi *et al*. 2006b). Gemin3/dp103/Ddx20 is a DEAD box RNA helicase (Charroux *et al*. 1999; Yan *et al*. 2003) that is essential for Sm core formation (Shpargel and Matera 2005) as well as for metazoan development (Mouillet *et al*. 2008; Shpargel *et al*. 2009). Gemin4 does not bind SMN directly, but is brought to the complex by virtue of its interaction with Gemin3 (Charroux *et al*. 2000; Otter *et al*. 2007). Gemin4 is necessary for Sm core formation *in vitro* (Shpargel and Matera 2005), but little else is known regarding the function of this protein.

Here, we show that *Gemin4* is an essential gene in the mouse and that the protein displays a dominant effect on the localization of SMN, Gemin3 and other members of the SMN complex when overexpressed in cultured human cells. This relocalization effect was completely dependent on the presence of an eight amino acid NLS within the N-terminal region of Gemin4 that is necessary and sufficient for targeting heterologous GFP constructs to the nucleus.

## Results

### Gemin4 contains a functional nuclear localization signal

Like most of the Gemin proteins, Gemin4 has no other known paralogs in the vertebrate proteome, and no known orthologs among non-vertebrate species (Kroiss *et al*. 2008). In humans and mice, Gemin4 is a 1058 aa protein with few known sequence features. Bioinformatic analysis using several web-based prediction tools suggested the existence of three classical (pat7 subtype) NLS sequences, one located within a slightly larger leucine zipper motif in the carboxy (C) terminal half and two others in the amino (N) terminal half of the protein (Fig. 1A).

**Figure 1.**
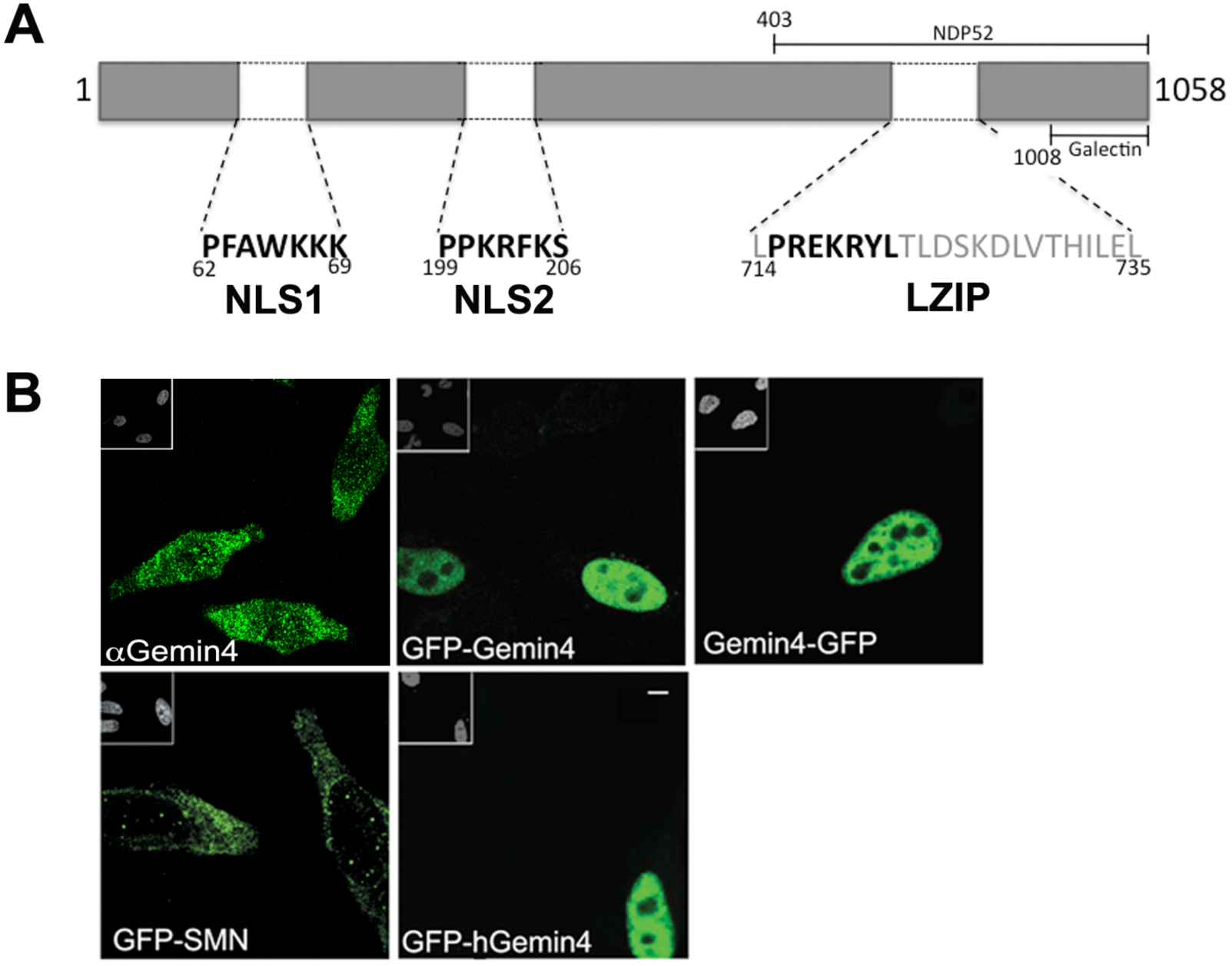
Gemin4 constructs and localization. (A) Schematic of Gemin4, illustrating structural features along with Galectin and NDP52 binding domains. Three predicted nuclear localization signal (NLS) sequences are shown (in bold type), one of which resides within a larger C-terminal leucine-rich motif (gray typeface). (B) Endogenous Gemin4 protein (α-Gemin4) localizes throughout the cytoplasm as well as in distinct nuclear foci called Cajal bodies. Full-length GFP-tagged mouse constructs (GFP-Gemin4 and Gemin4-GFP), as well as human Gemin4 (GFP-hGemin4) localize to the nucleoplasm when transiently transfected into HeLa cells. GFP-Smn localizes primarily to the cytoplasm, accumulating in nuclear foci, called Cajal bodies. Inserts in upper left corners are the DAPI stained nuclei. Scale bar, 5 *μ*m.

The sequences of human and mouse Gemin4 are greater than 84% identical and more than 90% similar. In order to distinguish endogenous from exogenous Gemin4 we created expression vectors that encode N- or C-terminal GFP-tagged versions of mouse Gemin4 (GFP-Gemin4 and Gemin4-GFP, respectively). As with SMN, endogenous Gemin4 typically localizes diffusely throughout the cytoplasm and in distinct nuclear foci called Cajal bodies (Charroux *et al*. 2000; Fig. 1B). Curiously, both myc and GFP N-terminally tagged mouse or human Gemin4 proteins predominantly localized to the nucleoplasm (Fig. 1B, and Fig. 4A,C). This accumulation in the nucleus was neither due to the placement of the tag nor due to the tag itself, as C-terminally tagged Gemin4-GFP (or -myc) showed the same pattern and GFP-SMN mirrored SMN’s endogenous distribution (Fig. 1B and data not shown). Thus, overexpression of Gemin4 results in a dominant gain-of-function phenotype, perhaps exposing a *cis*-acting NLS that is normally masked in the cytoplasm or titrating a *trans-acting* factor that normally facilitates Gemin4 nuclear export.

We mapped the NLS activity to the N-terminal half of Gemin4 using N- and C-terminal truncations (Fig. 2). Precise deletions of the predicted NLS motifs (Fig. 2A) identified an eight amino acid sequence that regulates GFP-Gemin4 nuclear import. Deletion of NLS1 (aa residues 62–69) or the leucine zipper (aa 714–735) had little effect, however, deletion of NLS2 (aa 199–206) relocalized the construct to the cytoplasm (Fig. 2B). Consistent with these findings, Lorson *et al*. (2008) reported that a ~50 aa sequence (aa 194–243) overlapping this region was required for Gemin4 import. The NLS2 motif is therefore necessary for nuclear import of Gemin4. To address whether NLS2 is sufficient for this process, we fused it to the carboxy terminus of GFP. Two constructs were expressed, one consisting of only the NLS2 sequence (NLS2min), and one with five additional residues flanking the NLS in both directions (NLS2ext). As shown in Fig 2B, native GFP displayed an overall pan-cellular localization pattern, whereas GFP-NLS2min was much more concentrated in the nucleus, albeit with a weak signal in the cytoplasm. In contrast, the GFP-NLS2ext construct was almost entirely nuclear (Fig. 2B). These observations demonstrate that the Gemin4 NLS2 motif is both necessary and sufficient for nuclear import.

**Figure 2.**
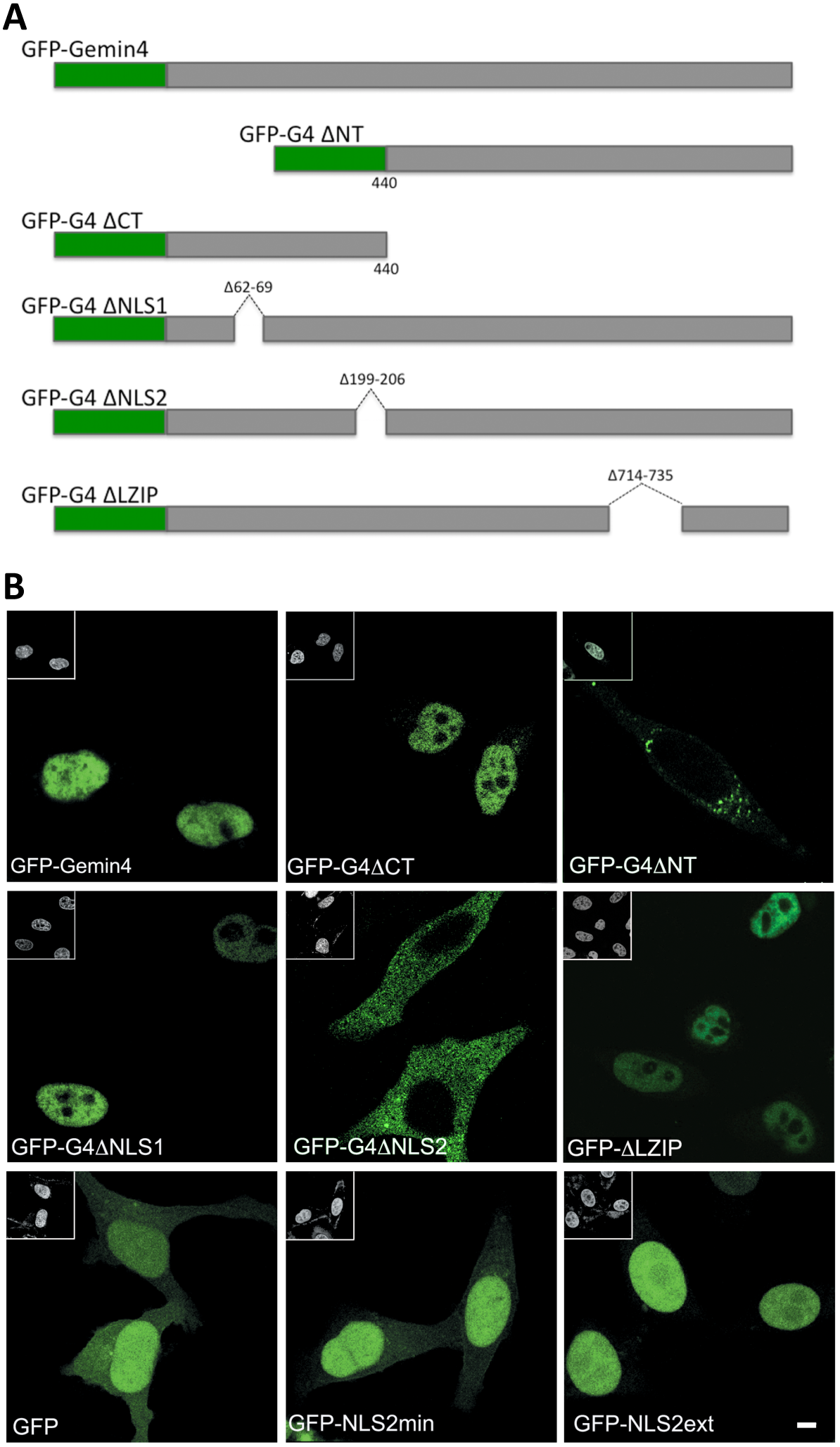
Identification of a functional NLS within mouse Gemin4. (A) Schematic of mouse Gemin4 protein. (B) (Top row) Full-length (GFP-Gemin4) and an N-terminal (GFP-G4∆CT) fragment of Gemin4 localize to the nucleoplasm. A C-terminal (GFP-+G4ANT) fragment accumulates in the cytoplasm. (Middle row) Expression of internal deletion constructs shows that only NLS2 is required for nuclear accumulation of Gemin4. (Bottom row) Although GFP-alone displays pan-cellular localization, the minimal NLS2 motif (GFP-NLS2min) is primarily nuclear. The extended NLS sequence (GFP-NLS2ext) is exclusively nuclear. Inserts in upper left corners of panels show the DAPI stained nuclei. Scale bar, 5 *μ*m.

### Gemin4 overexpression redistributes the SMN complex to the nucleoplasm

Although the tudor domain of SMN can interact directly with Importin-β (Narayanan *et al*. 2004), Gemin4 is the only SMN complex member that contains an NLS. We therefore examined the ability of Gemin4 to influence the subcellular localization of other SMN complex proteins. HeLa cells were transiently transfected with GFP-Gemin4 and immunostained for SMN, Gemin2, Gemin3 and Unrip (Fig. 3). In each case, overexpression of GFP-Gemin4 caused relocalization of the endogenous protein to the nucleus. There was a clear dosage effect to the redistribution. In weakly transfected cells, SMN was qualitatively reduced (but still visible) in the cytoplasm (Fig. 3A,B). In cells strongly expressing GFP-Gemin4, endogenous SMN was no longer detectable in the cytoplasm and the nuclear SMN foci (Cajal bodies) were much less frequent (Fig. 3A,B). Similar effects on the cellular distributions of SMN and Gemin2 were observed upon overexpression of myc-Gemin4 (Fig. 4A,C). Quantitative analysis showed that expression of myc- or GFP-Gemin4 significantly relocalized endogenous SMN and Gemin2 to the nucleoplasm (Fig. 4B,F).

**Figure 3.**
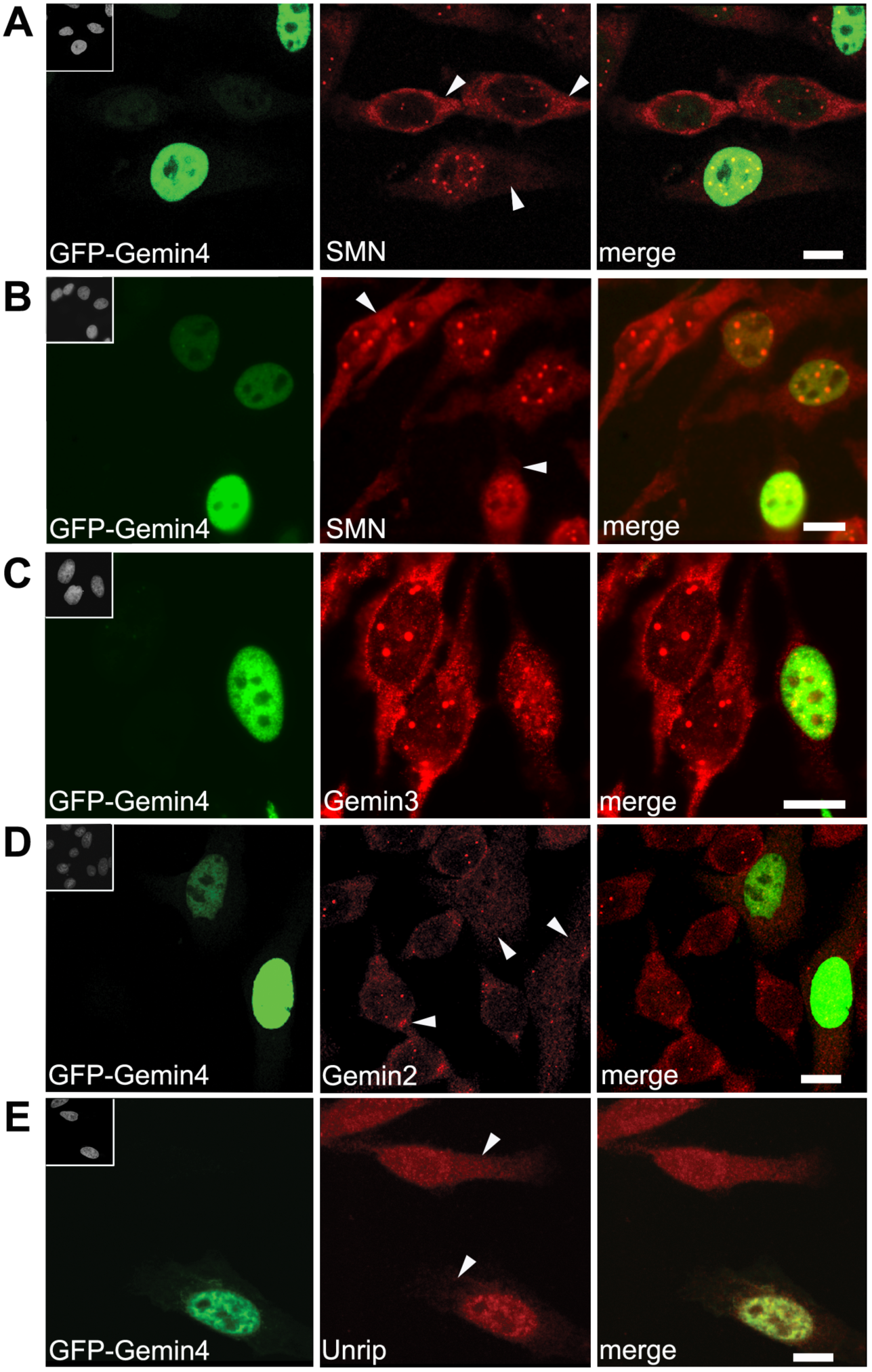
Overexpression of GFP-Gemin4 causes mislocalization of SMN and other members of the SMN complex to the nucleoplasm. HeLa cells were transfected with GFP-Gemin4 (shown in green) and then co-stained (in red) with antibodies targeting SMN (A and B), Gemin3 (C), Gemin2 (D), or Unrip (E). Arrowheads are used to illustrate the reduced cytoplasmic staining of endogenous proteins in the transfected cells compared to the untransfected cells. Inserts in upper left corners of the GFP images show the DAPI stained nuclei. Scale bars, 10 *μ*m.

**Figure 4.**
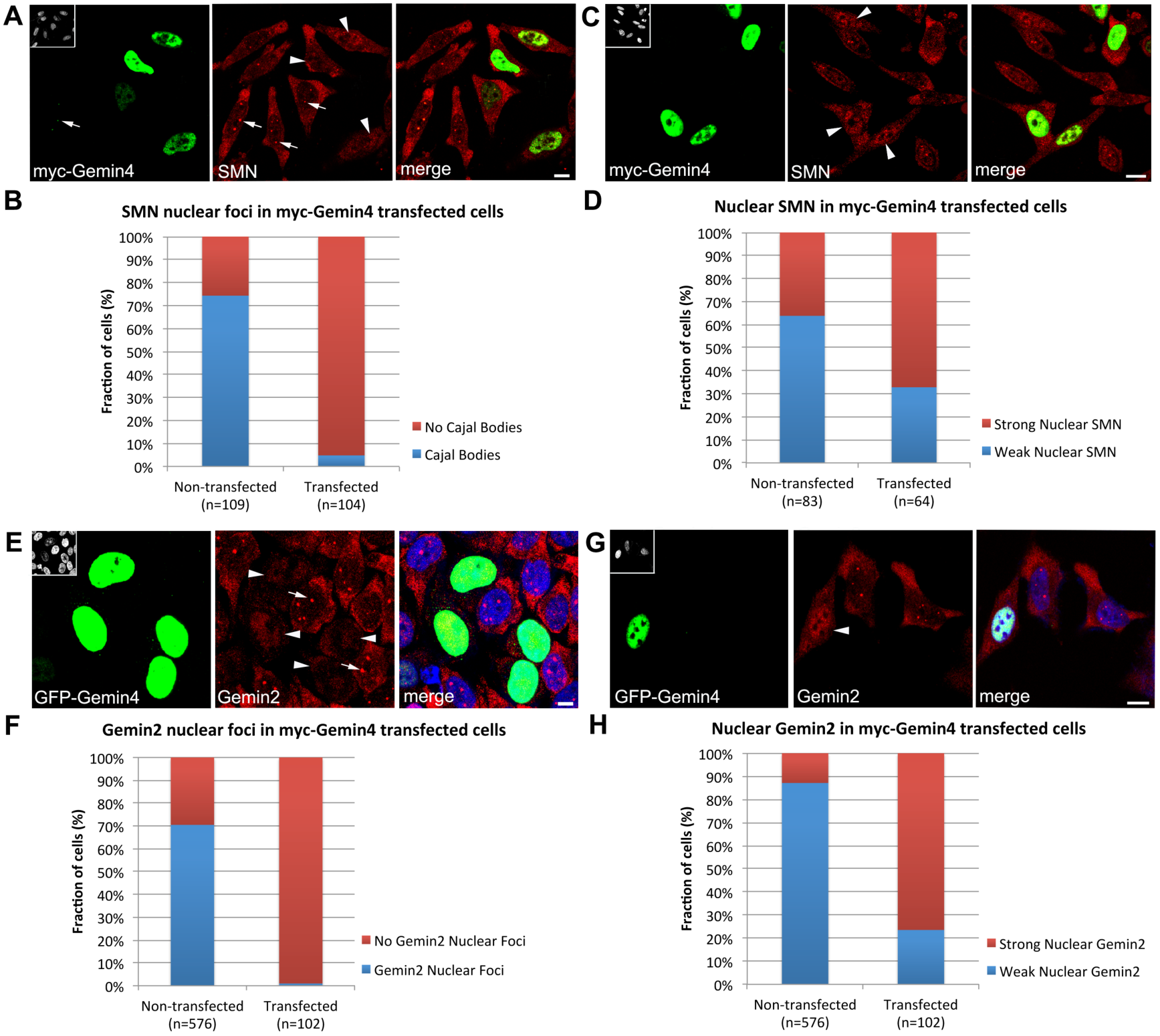
Quantification of SMN and Gemin2 phenotypes in HeLa cells overexpressing Gemin4. HeLa cells were transfected with either myc- (panels A-D) or GFP-Gemin4 (E-H) and then co-stained with antibodies targeting SMN or Gemin2, as shown. Each nucleus was scored for the presence or absence of nuclear foci (B and F) as well as for nucleoplasmic accumulation (D and H). Scale bars, 5 *μ*m.

### High levels of Gemin4 overexpression disrupts Cajal bodies

Gemin4 overexpression drives cytoplasmic SMN complexes into the nucleoplasm, but these proteins frequently fail to concentrate in Cajal bodies (Fig. 3). We quantified the loss of SMN, Gemin2, and Gemin3 nuclear foci in myc- or GFP-Gemin4 expressing cells and found that the ability of these SMN complex components to accumulate within nuclear foci was significantly inhibited (Fig. 4 and data not shown). To determine if the loss of these nuclear foci in myc- or GFP-Gemin4 expressing cells was due to Cajal body disassembly, we examined the distribution of known Cajal body marker proteins: coilin, WDR79 and NPAT (Fig. 5A-C). Nuclear foci with these Cajal body markers were largely unaffected in cells expressing lower levels of GFP-Gemin4, but we observed an appreciable reduction of nuclear foci in cells with high GFP-Gemin4 expression (Fig. 5C). We then quantified the loss of coilin-positive nuclear foci and found a significant increase in cells that contained no distinguishable nuclear foci (Fig. 5D). Thus, the observed Gemin4-dependent relocalization of SMN and its binding partners to the nucleoplasm perturbs Cajal body integrity in a dose dependent manner. Because coilin interacts with SMN and mediates recruitment of the SMN complex to Cajal bodies (Hebert *et al*. 2002; Hebert *et al*. 2001), it is likely that the SMN binding sites on coilin are simply swamped by the preponderance of nuclear SMN.

**Figure 5.**
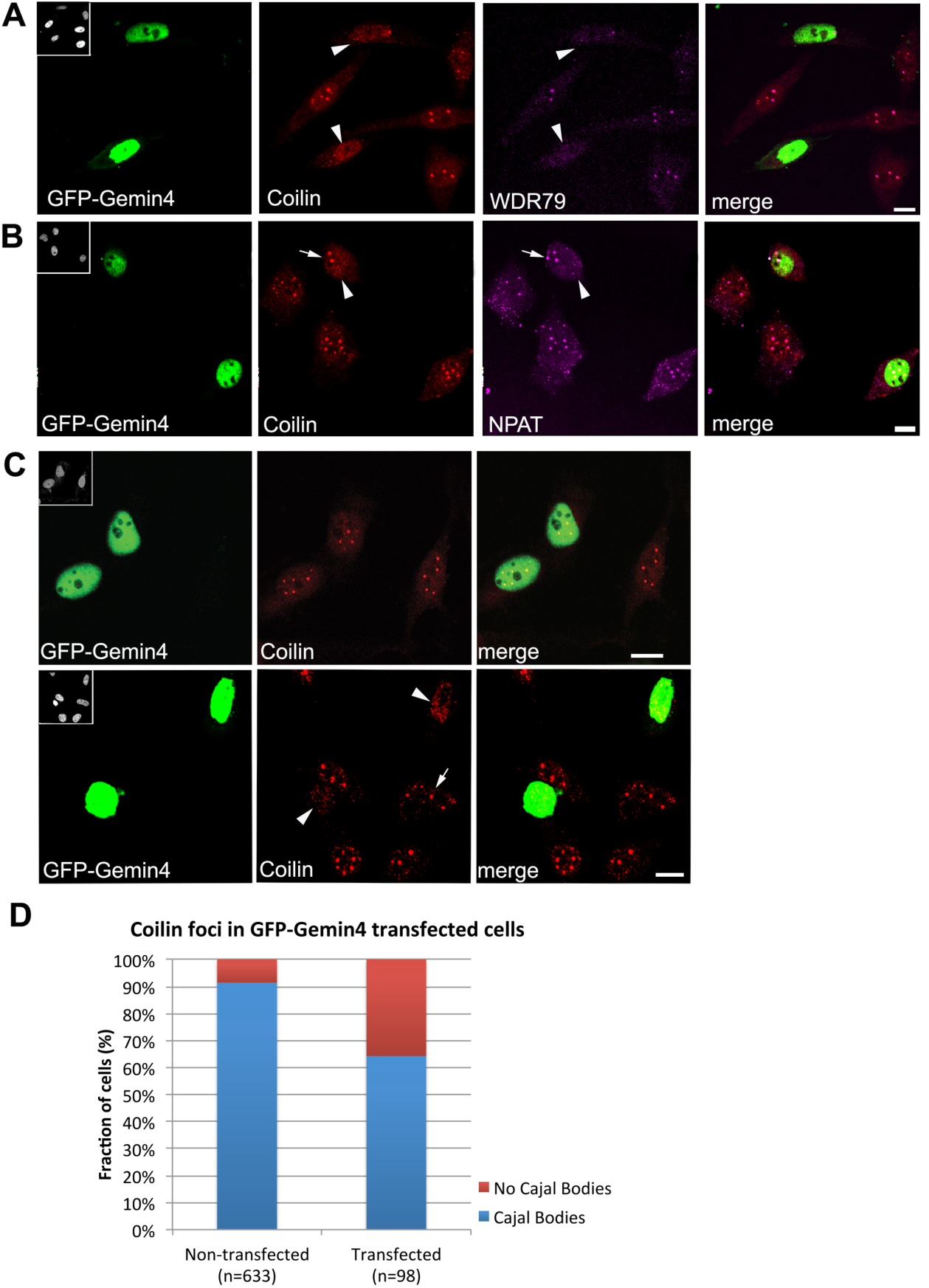
Cajal bodies are disrupted by high levels of GFP-Gemin4 expression. HeLa cells were transfected with GFP-Gemin4 (shown in green) and then co-stained (in red) with antibodies targeting Coilin (A-C), WDR79 (A) and NPAT (B). Inserts in upper left corners of GFP images show the DAPI stained nuclei. Arrowheads are used to identify transfected cells. Arrows indicate coilin foci corresponding to Cajal bodies. Cells strongly expressing GFP-Gemin4 show major disruptions to Cajal bodies, whereas lower expression levels had little effect (B). Nuclei were scored for the presence or absence of coilin foci (D). Chi squared analysis reveals a significant difference between the two cell populations, p << 0.001. Scale bars, 10 *μ*m.

### The Gemin4 NLS is required for SMN and Cajal body disruption

To further examine the critical Gemin4 regions involved in SMN relocalization, we overexpressed Gemin4 lacking the NLS (myc-G4∆NLS2). We predicted that NLS2 was responsible for the disruption of SMN nuclear foci upon overexpression of Gemin4. We found that SMN nuclear foci were relatively unaffected by myc- or GFP-Gemin4∆NLS2 expression (Fig. 6A,B), demonstrating that the loss of SMN nuclear foci upon Gemin4 overexpression requires the presence of Gemin4 NLS2. In contrast, both Gemin2 and Gemin3 nuclear foci were significantly reduced upon myc- or GFP-Gemin4∆NLS2 expression (Fig. 6D,F). This observation indicates that Gemin2 and Gemin3 are sequestered by Gemin4∆NLS2 in the cytoplasm. Notably, the reduction in Gemin3 nuclear foci was much more robust than that of Gemin2 (Fig. 6D,F). This is perhaps due to the fact that Gemin4 is a direct binding partner of Gemin3, whereas Gemin2 is not (Otter *et al*. 2007). Additionally, we note that Coilin nuclear foci were observed in nearly all myc- or GFP-Gemin4∆NLS2 transfected and control cells (Fig. 6G,H). Taken together, these data suggest that the disruption of coilin and SMN nuclear foci depends upon a functional Gemin4 NLS.

**Figure 6.**
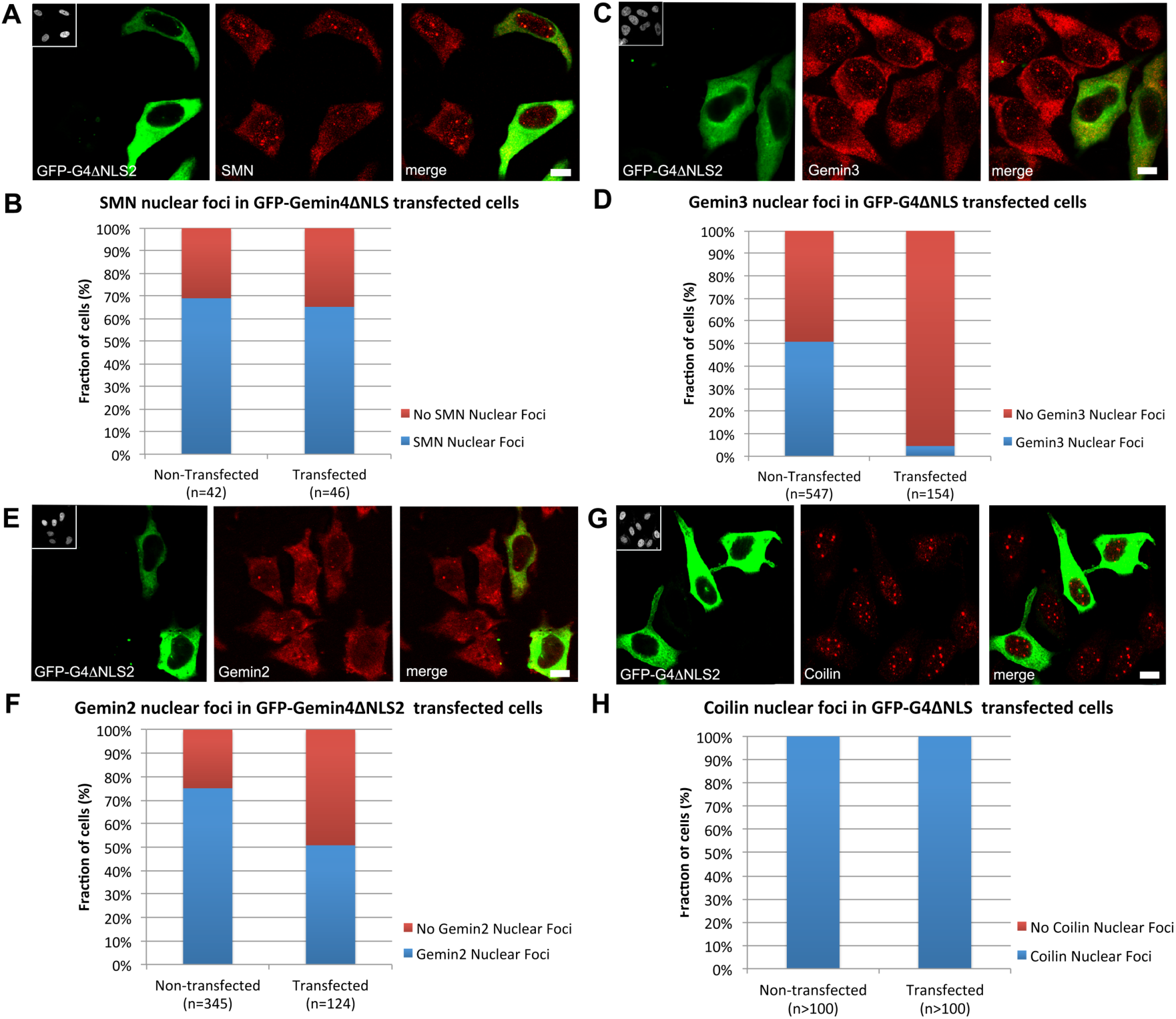
Immunofluorescence of endogenous SMN complex and coilin proteins in cells transfected with GFP-Gemin4∆NLS2. HeLa cells were transfected with GFP- Gemin4∆NLS2 (shown in green) and then co-stained (in red) with antibodies targeting SMN (A), Gemin2 (C), Gemin3 (E) or Coilin (G). As quantified above, GFP- Gemin4∆NLS2 expression disrupts Gemin2 (F) and Gemin3 (D), but not SMN (B) or Coilin (H) nuclear foci. Chi squared analysis shows p=1.0 for coilin, p>0.7 for SMN and p << 0.001 for Gemin3, and Gemin2. Inserts in upper left corners of GFP images are the DAPI stained nuclei. Arrowheads are used to identify transfected cells with reduced cytoplasmic staining of endogenous proteins. Arrows indicate nuclear foci in non-transfected cells. Scale bars, 5 *μ*m.

### Nucleoplasmic relocalization of Gemin3 requires a C-terminal motif in Gemin4

As mentioned above, Gemin4 does not interact directly with SMN; it is thought to be tethered to the SMN complex via its strong and direct interaction with Gemin3 (Charroux *et al*. 2000; Otter *et al*. 2007). As shown in Figs. 3C and 7A, overexpression of GFP-Gemin4 mislocalizes endogenous Gemin3 to the nucleoplasm. We used this effect as a way to map the interaction between Gemin4 and Gemin3. Interestingly, the GFP-Gemin4ΔCT construct, which contains the NLS and localizes to the nucleus, did not perturb localization of endogenous Gemin3 (Fig. 7B). This finding suggests that the truncated Gemin4 no longer interacts with Gemin3 and that the domain responsible for this interaction is located in the C-terminal half of Gemin4. Consistent with this interpretation, we found that relocalization of Gemin3 depended on the presence of the putative leucine zipper motif in Gemin4. Cells transfected with GFP-Gemin4∆zip displayed a Gemin3 localization pattern comparable to that of non-transfected cells (Fig. 7C). These data demonstrate that the Gemin4-dependent translocation of the SMN complex to the nucleus depends on a leucine-rich domain in the Gemin4 C-terminus and further suggest that Gemin4 may bind to Gemin3 via this motif.

**Figure 7.**
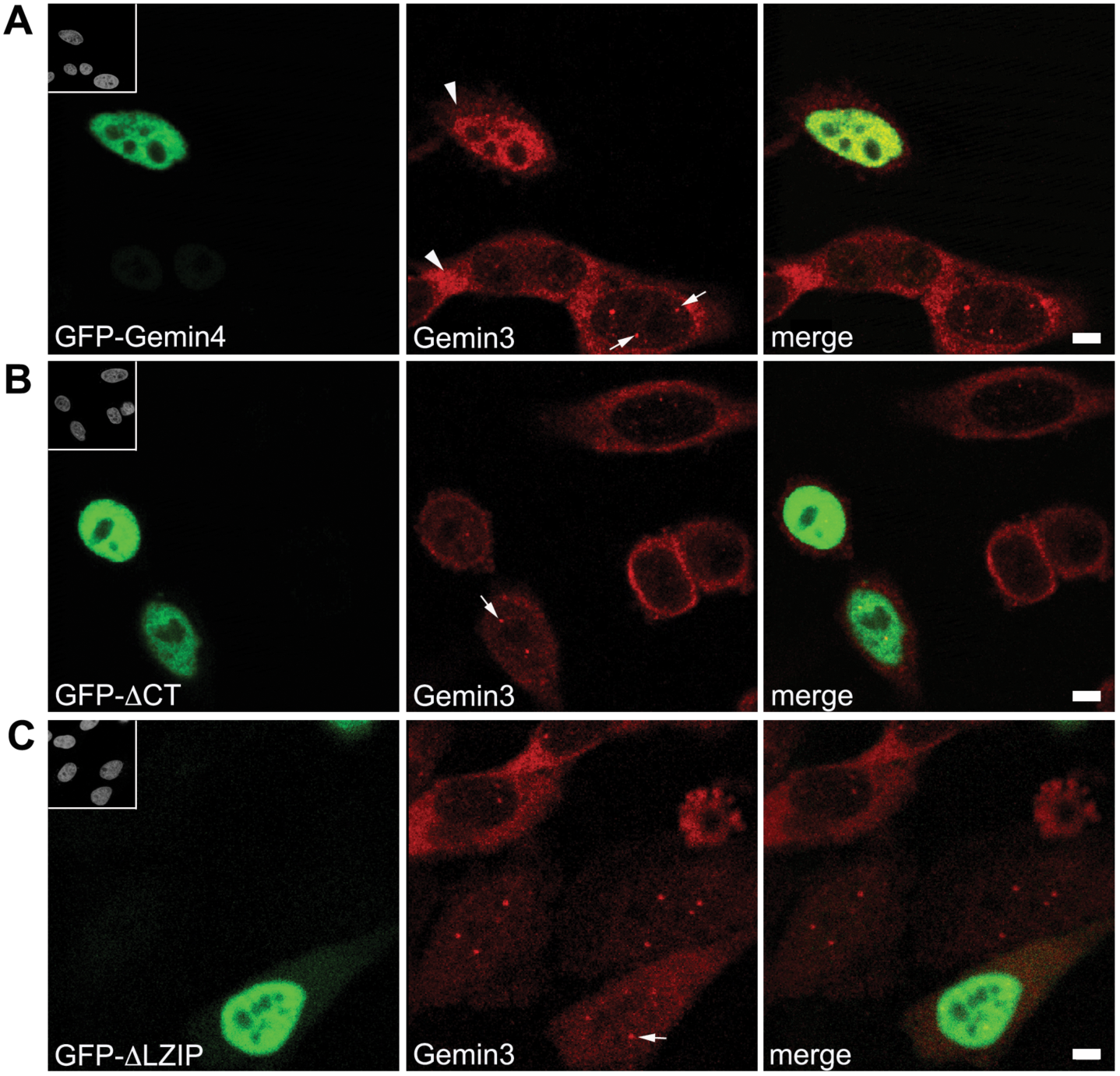
Gemin4-mediated nuclear relocalization of Gemin3 requires a C-terminal region within Gemin4. HeLa cells were transfected with various GFP-Gemin4 constructs (green) and then co-stained using anti-Gemin3 (red). Expression of full-length GFP-Gemin4 (A) relocalized Gemin3 to the nucleoplasm, whereas expression of an N-terminal truncation, GFP-ΔCT (B), or an internal deletion, GFP-ΔLZIP (C), did not. Arrowheads in panel (A) are used to illustrate the reduced cytoplasmic staining of endogenous Gemin3 in transfected cells, as compared to untransfected cells. Arrows mark Cajal bodies. Scale bars, 5 *μ*m.

### Characterization of a Gemin4 gene trap allele in the mouse

Experiments in cultured cells can contribute a great deal of information for studying protein function. However, it is difficult to extrapolate the relevance of a given protein from single cell studies to the importance that it may have in the context of organismal development. Therefore, we created and characterized a loss-of-function *Gemin4* mouse model. We obtained mouse embryonic stem (ES) cells (Lexicon Genetics) that contain a retroviral insertion (Zambrowicz *et al*. 1998) in the single *Gemin4* intron. This “gene trap” insertion is derived from an engineered retroviral cassette that preferentially inserts itself into upstream introns within the mouse genome. The genomic organization of *Gemin4* is ideal for this kind of gene-trapping scheme because it is a single-intron gene and the upstream exon (exon 1) is particularly short, encoding only the first three aa residues (see Fig. 8). Exon 2 thus contains essentially the entire protein coding sequence. The gene-trap contains an upstream element consisting of a 5’ splice acceptor site fused to a β-galactosidase/neomycin-resistance (β-geo) cassette that also contains a transcription termination sequence. The β-geo cassette is transcribed from the endogenous promoter of the *Gemin4* gene, located on mouse chromosome 11. The result is a fusion transcript in which the exon upstream of the insertion site is spliced in-frame to the β-geo reporter. The fusion transcript encodes three residues of the Gemin4 polypeptide fused to the β-geo cassette. The downstream element contains a puromycin-resistance cassette along with its own PGK promoter that ‘hijacks’ expression of exon 2. The gene trap has stop codons introduced in all three downstream reading frames. There is a splice donor site that fuses this transcript with exon 2 (Fig. 8A).

**Figure 8.**
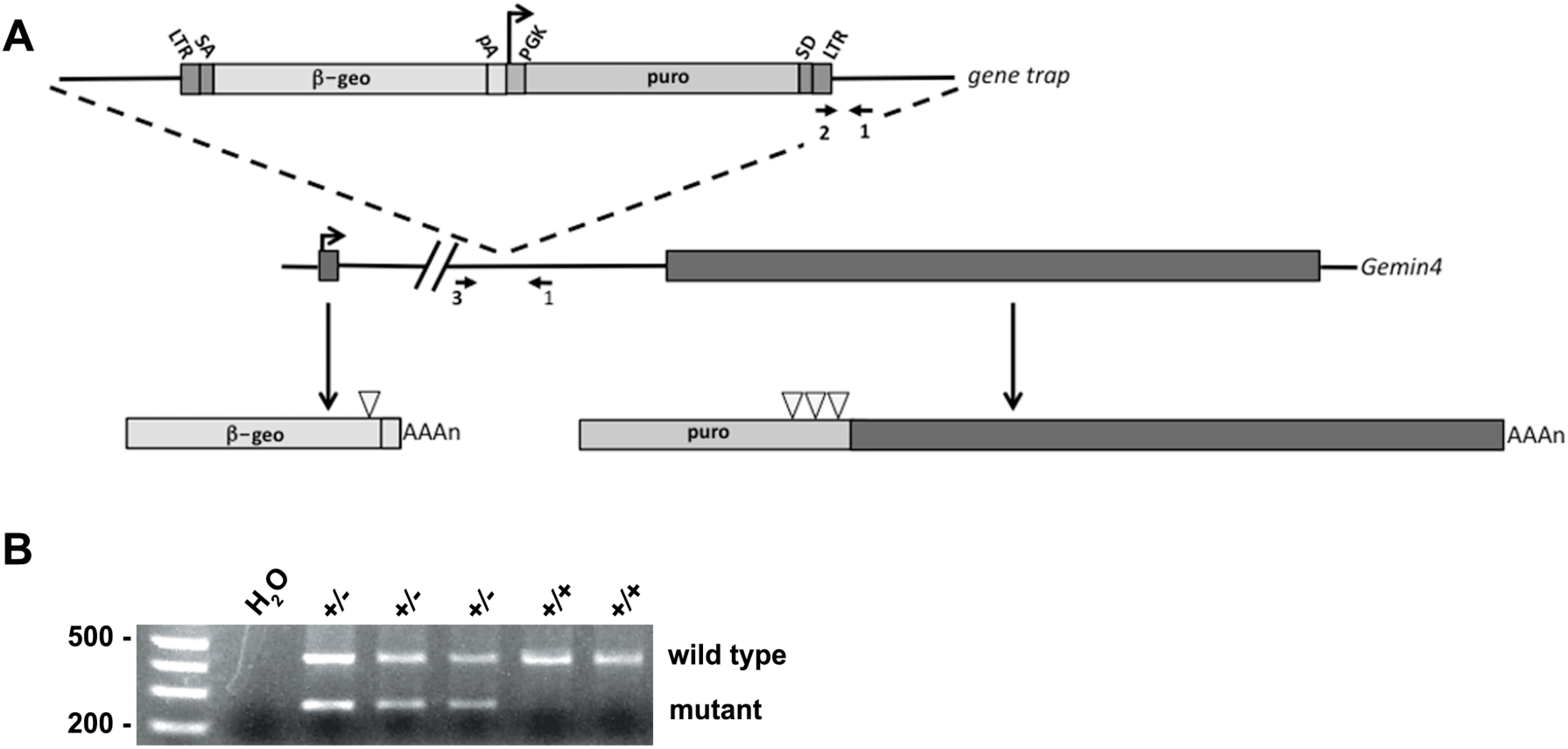
Schematic of Gemin4 gene trap. (A) Retroviral insertion of the gene trap vector into the single intron of murine *Gemin4* creates an upstream transcript that incorporates exon 1 with a β-geo cassette. Exon 2 is thus trapped into expressing a puromycin cassette that contains stop codons (open triangles) introduced in all three reading frames just upstream of the exonic sequence. The locations of the three PCR primers used in the genotyping assay are shown (arrows). LTR, retroviral long terminal repeat; SA, splice acceptor; SD, splice donor; PGK, phosphoglycerate kinase promoter region. (B) Germline transmission was observed in two of four chimeric males when crossed with a wild-type female. Primers 1 and 2 (shown in panel A) generate a 250 bp band derived from the mutant chromosome, whereas primers 1 and 3 generate a 450 bp wild-type band.

Blastocysts harboring the mutant ES cells were injected into pseudopregnant females, resulting in four chimeric males. Two of these mice displayed germline transmission of the gene-trap, which was ascertained by PCR genotyping of the founder progeny (Fig. 8B). Founder mice were then backcrossed for three generations onto the C57BL/6J genetic background.

### Gemin4 is an essential gene in the mouse

Heterozygous animals were intercrossed and the resulting offspring were PCR genotyped at various developmental stages. Whereas heterozygous and wild-type progeny were readily detected at postnatal day 1 (P1), embryonic day 13.5 (E13.5), E7.5 and E5.5, no *Gemin4* homozygotes were detected at any of these time points (Table 1 and data not shown). These findings demonstrate that although *Gemin4* is thought to be an evolutionary newcomer to the SMN complex (it has been identified only in vertebrate genomes; Kroiss *et al*. 2008), it is an essential mammalian gene. The data are also consistent with gene targeting experiments showing that mouse *Gemin2* and *Gemin3* knockouts are also early embryonic lethal mutations (Jablonka *et al*. 2002; Mouillet *et al*. 2008).

**Table 1.** *Gemin4* is an essential gene in the mouse. *Gemin4* heterozygotes were intercrossed and the genotypes of F1 progeny were analyzed by PCR at P1, E13.5 and E7.5.

|  | *Gemin4*<sup>+/+</sup> | *Gemin4*<sup>+/-</sup> | *Gemin4*<sup>-/-</sup> | Total |
| --- | --- | --- | --- | --- |
| P1 | 30 | 69 | 0 | 99 |
| E13.5 | 16 | 36 | 0 | 52 |
| E7.5 | 18 | 33 | 0 | 51 |

### Transgenic suppression of Smn lethality by SMN2 is background-dependent

The human genome contains two copies of the SMN gene, *SMN1* and *SMN2*. Homozygous loss of *SMN2* is asymptomatic, but loss of SMN1 results in a neuromuscular disease, Spinal Muscular Atrophy (SMA) (Lefebvre *et al*. 1995). SMA is caused by hypomorphic reduction of SMN protein levels, whereas complete loss of gene function is embryonic lethal (reviewed in Burghes and Beattie 2009; Cauchi 2010). The mouse genome contains only a single copy of the *Smn* gene, null mutation of which is early embryonic lethal (Schrank *et al*. 1997). Transgenic expression of human *SMN2* in the *Smn* knockout background rescues embryonic lethality, and recapitulates the SMA type I phenotype (Hsieh-Li *et al*. 2000; Monani *et al*. 2000).

In order to analyze the effect of *Gemin4* copy number on the SMA phenotype, we generated a colony of *Smn^+/-^;SMN2^+/+^* mice and backcrossed them onto the C57BL/6J inbred background for >6 generations. We then intercrossed the animals and were surprised to find that the *Smn^-/-^;SMN2^+/+^* genotype was never detected postnatally (Table 2). Similar results were observed on a mixed C57BL/6J;129X1/SvJ background (Table 2). In contrast, and consistent with previous findings (Monani *et al*. 2000), *Smn^-/-^;SMN2^+/+^* animals were detected in their expected numbers on the FVB/NJ hybrid background. The expected 1:2 ratios of wild-type to heterozygous mice in the non-FVB strains (Table 2) suggest that the *Smn^-/-^;SMN2^+/+^* embryos in those strains died *in utero*. We conclude that *SMN2’s* function as a genetic suppressor of the *Smn* embryonic lethal phenotype is background dependent.

**Table 2.** Analysis of genetic background contributions to the SMA phenotype. C57BL/6J and FVB/NJ mice that had been bred on pure (backcrossed more than 10 generations) inbred strains or those from a mixed background (C57BL/6J;129X1/SvJ).

|  | *Smn*<sup>+/+</sup>,<br>*SMN2*<sup>+/+</sup> | *Smn*<sup>+/-</sup>,<br>*SMN2*<sup>+/+</sup> | *Smn*<sup>-/-</sup>,<br>*SMN2*<sup>+/+</sup> | Total |
| --- | --- | --- | --- | --- |
| C57BL/6J;129X1SvJ | 19 | 42 | 0 | 61 |
| C57BL/6J | 20 | 33 | 0 | 53 |
| FVB/NJ | 20 | 40 | 28 | 88 |

### Gemin4 does not function as a genetic modifier of mouse Smn

In order to assay the effects of *Gemin4* copy number on the SMA phenotype, we crossed the *Gemin4* gene trap onto the SMA background to obtain *Gemin4^+/-^;Smn^+/-^;SMN2^+/+^* animals. We intercrossed these mice and analyzed their progeny. As shown in Table 3, SMA mice were present in expected numbers irrespective of the *Gemin4* copy number. This failure to exacerbate (or ameliorate) the early lethality phenotype was puzzling, considering the fact that both *Gemin2* and *Gemin3* heterozygous mice produce roughly half the protein levels compared to wild-type and have associated phenotypes as a result (Jablonka *et al*. 2002; Mouillet *et al*. 2008). Notably, *Gemin4* heterozygotes are phenotypically indistinguishable from their wild-type littermates.

**Table 3.** *Gemin4* heterozygotes do not modify the early lethality phenotype of type I SMA animals. Parental mice with the genotype of *Gemin4^+/-^;Smn^+/-^;SMN2^+/+^* were intercrossed and the resulting progeny were PCR genotyped to look for SMA type I-like mice that were either wild type or heterozygous for *Gemin4*.

|  | Observed | Expected |  |
| --- | --- | --- | --- |
| *Gemin4*<sup>+/+</sup>; *Smn*<sup>-/-</sup>; *SMN2*<sup>+/+</sup> | 9 | 10 |  |
| *Gemin4*<sup>+/-</sup>; *Smn*<sup>-/-</sup>; *SMN2*<sup>+/+</sup> | 22 | 20 | p-value = 0.859 |

We therefore examined the expression profile of Gemin4 in various tissues. Consistent with microarray expression profiles (NCBI gene ontology database profile GDS3142), Gemin4 was readily detectable in all the tissues we examined (data not shown). Although wild-type adult and neonatal mice expressed approximately two-fold greater amounts *Gemin4* mRNA than their heterozygous littermates, the protein levels from these animals were consistently equivalent (Fig. S1). It is possible that post-translational mechanisms exist to regulate Gemin4 protein levels in heterozygotes. We screened numerous commercial and non-commercial antibodies against Gemin4 for their ability to recognize the mouse protein (not shown). Although a few worked nominally well for westerns, unfortunately none of them were suitable for immunofluorescence experiments. Thus we were unable to analyze *Gemin4* preimplantation embryos to carry out additional phenotypic analyses. We also note that *Gemin4* heterozygous mice are phenotypically identical to wild-type mice and, as a result, they are unable to modify the SMA phenotype.

## Discussion

Gemin4 contains a functional NLS in its N-terminal domain that is necessary and sufficient for nuclear import of exogenous cargoes. Curiously, the localization pattern of epitope-tagged versions of Gemin4 does not mirror that of the endogenous protein. Ectopically expressed Gemin4 is primarily nucleoplasmic, whereas the endogenous protein is localized throughout the cytoplasm, and in nuclear Cajal bodies. Mislocalization of the tagged constructs was not due to the presence of the tag itself, as a construct containing a deletion of the NLS (GFP-Gemin4∆NLS2; Fig. 2) was completely excluded from the nucleus, with the exception of a very small number of cells that had distinct myc-Gemin4∆NLS2 nuclear foci that colocalized with SMN (Fig. 6A). This rare targeting to Cajal bodies suggests that a small fraction of myc-Gemin4∆NLS2 proteins may gain access to the nucleus by piggybacking onto Gemin3 and the rest of the SMN complex. We also discovered that relocalization of Gemin3 depends on the presence of the leucine zipper motif in Gemin4 (Fig. 7). Because Gemin4∆NLS2 retains this motif, it is somewhat surprising to find that Gemin4∆NLS2 is apparently limited in its ability to piggyback onto Gemin3 for import into the nucleus. The fact that SMN, but not other SMN complex components like Gemin2 and Gemin3, retains the ability to localize to Cajal bodies upon overexpression of Gemin4∆NLS2 suggests that SMN can be imported independently from the other Gemins. Consistent with this observation is the fact that SMN was shown to directly interact with Importin-β (Narayanan *et al*. 2004; Narayanan *et al*. 2002).

Gemin4 is thought to be tethered to the SMN complex via its interaction with Gemin3, and it was shown to form a subcomplex together with Gemin5 to create a Gemin3/4/5 heterotrimer (Battle *et al*. 2007; Carissimi *et al*. 2005; Carissimi *et al*. 2006b). Overexpression of Gemin4 likely alters the stoichiometry of the various members of the SMN complex (or subcomplexes thereof). Thus it is also a bit surprising to find that Gemin4 overexpression results in the nucleoplasmic accumulation of SMN and all other tested members of the complex. These findings indicate that Gemin4 may play a regulatory role involved in the nuclear import of the SMN complex. SMN enters the nucleus during import of newly-formed snRNPs and is part of a two-component nuclear import signal (Fischer *et al*. 1993; Narayanan *et al*. 2004; Narayanan *et al*. 2002). Depending on the combination of binding partners in a given complex or subcomplex, it is possible that the Gemin4 NLS is masked. Under certain conditions, the Gemin4 NLS might serve to tip the balance of import in one direction or the other.

### Characterization of Gemin4 loss-of-function mice

Our data clearly demonstrate that *Gemin4* is an essential mammalian gene. Importantly, null mutations in *Smn, Gemin2* and *Gemin3* are all embryonic lethal, demonstrating that these genes are also essential (Jablonka *et al*. 2002; Mouillet *et al*. 2008; Schrank *et al*. 1997). U snRNP biogenesis is an essential cellular process that ensures the availability of splicing factors required for gene expression. Previous knockdown experiments revealed that depletion of Gemin4 (or Gemin3) causes an intermediate, yet significant, loss of U snRNP assembly activity in a HeLa cells (Shpargel and Matera 2005). This intermediate effect was not overly detrimental, as cell death was not nearly as pronounced as it is upon SMN knockdown (Lemm *et al*. 2006, and our unpublished observations). These observations suggest that Gemin4 and Gemin3 may be dispensable when it comes to the basal levels of U snRNPs needed to maintain cells in culture. However, during mammalian development U snRNA and snRNP levels increase dramatically from the 2–16 cell stage to the blastocyst stage (Dean *et al*. 1989; Lobo *et al*. 1988). The SMN complex would most likely need to operate at peak efficiency to account for the large increase in U snRNP production during these critical stages of development and the full complement of Gemins may be required to stabilize the complex or directly aid in assembly of snRNPs (Strzelecka *et al*. 2010). This idea is bolstered by the fact that *Smn, Gemin2, Gemin3* and *Gemin4* null mice all die during early embryogenesis (Jablonka *et al*. 2002; Mouillet *et al*. 2008; Schrank *et al*. 1997; this work).

Gemin4 was also reported to reside in certain mammalian miRNP complexes (Dostie *et al*. 2003; Hutvagner and Zamore 2002; Mourelatos *et al*. 2002). Given their wide range of gene regulatory roles, including post-transcriptional mRNA cleavage or translational repression (He and Hannon 2004), miRNPs are required for mammalian development (Bernstein *et al*. 2003; Chen *et al*. 2004; Houbaviy *et al*. 2003). Gemins 3 and 4 were reported to bind in a separate complex along with the Argonaute protein, eIF2C2/Ago2 (Mourelatos *et al*. 2002; Nelson *et al*. 2004), although the significance of these findings has not been explored. Because relatively few proteins make up this miRNP subclass, it is likely that removal of any of its members would render it non-functional. Gemin4 is not known to have a paralog or redundant equivalent; it is reasonable to assume that if Gemin4 is required for miRNP assembly, loss of its function would be detrimental to this pathway as well as for snRNP biogenesis.

In summary, we have shown that Gemin4 has the potential to redirect the localization of other SMN complex members from the cytoplasm to the nucleoplasm in cultured mammalian cells. Gene disruption in mice demonstrated that *Gemin4* is required for embryonic viability. We also found that *SMN2* failed to rescue the embryonic lethality phenotype of *Smn* knockouts, when bred on the C57BL/6J inbred background. These data demonstrate the existence of additional genetic modifiers of SMA. We were unable to conclude whether *Gemin4* is such a modifier, as we found that heterozygous *Gemin4* mice express wild-type levels of protein.

## Materials and Methods

### Plasmid construction

Mouse total brain cDNA was used to PCR amplify mouse Gemin4. The amplicon was cloned into the pEGFP (Clontech) or pcDNA-myc (Invitrogen) vectors to express both N- and C-terminally tagged proteins. The QuickChange site-directed mutagenesis kit (Stratagene) was used to create the deletion constructs following the manufacturer’s protocol (primer sequences available upon request).

### Immunofluorescence microscopy

HeLa cells were seeded on two-well glass slides and grown in an incubator at 37 °C with 5% CO_2_ in DMEM (Mediatech) supplement with 10% BSA and 1% Penicillin/Streptomycin. HeLa cells were grown until they reached the 60% confluency, and transiently transfected using Effectene transfection kit (Qiagen, manufacturer protocol). Cells were harvested 24 h later, fixed in 4 % PFA in 1x PBS solution for 20 min at RT, and permeabilized in Triton X-100 followed by 3 washes in 1x PBS for 5 min at RT. Immunofluorescence experiments were performed by incubations of primary antibody diluted in PBS containing 3% BSA followed by incubation with secondary Ab. Antibodies used were as follows: anti-SMN mAb (clone 8, BD biosciences, 1:200), anti-dp103/Gemin3 mAb (4G7, 1:10), anti-Gemin4 mAb (clone 3E1, Abnova, 1:10), anti-WDR79 mAb (ab77333, abcam, 1:100), anti-NPAT mAb (DH4, gift from J. Zhao, 1:50), anti-coilin pAb (R124, 1:400), anti-Unrip mAb (3G6, 1:10), anti-Gemin2 mAb (6084–100 abcam 1:200), Alexa 594 goat anti-mouse and goat anti-rabbit secondary antibody (Invitrogen) were used. The incubations were carried out at 37 °C for an hour, stained 3 min with DAPI, washed with 1x PBS, and covered with antifade solution to avoid bleaching. Laser confocal fluorescence microscopy was performed with Leica TCS SP5 high speed and high-resolution spectral confocal microscope. Images from each channel were taken within a single plane with an objective of 63x with a 3x zoom factor and recorded separately and the files were merged as needed.

### Mouse lines and crosses

Various inbred mouse strains used in this study (FVB/NJ, C57BL/6J and 129Sv/J) were obtained from the Jackson Laboratory. Wild-type animals thus obtained were used for colony maintenance and to outcross *Gemin4* and *Smn* mutants onto the various genetic backgrounds described in the results section. Mice carrying a human *SMN2* transgene in the background of a null mutation in the endogenous *Smn* gene (SMA type I mice: FVB.Cg-Tg(*SMN2*)89*^Ahmb^*;*Smn^tm1Msd^*/J) were obtained from the Jackson Laboratory. *Gemin4* mice were created by purchasing ES cells with the *Gemin4* gene-trap cassette from Lexicon Genetics and these cells were injected into donor blastocysts and subsequently injected into a pseudopregnant female mouse by the Case Western Reserve University transgenic mouse facility. All strains were maintained on a standard diet of 50/10 food pellets and sterile water. These mice were housed in micro-isolation chambers. Breeding pairs for SMA type I mice consisted of mice that were homozygous for the transgene and heterozygous for the knockout allele, which resulted in pups that display the SMA phenotype and control littermates. Breeding pairs for *Gemin4* mice were heterozygous for the gene trap cassette. All mice were humanely euthanized according to protocols and standards set forth by the appropriate Institutional Animal Care and Use Committees (IACUC): the Case Western Reserve University Animal Resource Center (CWRU ARC) and the University of North Carolina Division of Laboratory Animal Medicine (UNC DLAM).

### Reverse transcription and quantitative real-time PCR

RNA was extracted from homogenized liver using the RNeasy kit (Qiagen; manufacturer’s protocol), including an RNase-free DNase (Qiagen) on-column digestion step to remove genomic DNA. The SuperScript First Strand Synthesis System for RT-PCR (Invitrogen) was used to synthesize cDNA in a 20 *μ*l reaction containing 5 *μ*g RNA, 179 ng random hexamer primer and 40 U RNaseOut RNase inhibitor (Invitrogen). The reverse transcription reaction was performed (Invitrogen SuperScript; manufacture’s protocol). In addition to the PCR mastermix buffer (gotaq, Promega), each PCR reaction mixture (20 *μ*l) contained 1 *μ*l of cDNA 1:10 dilution. The Primers used are listed below. PCR conditions consisted of one step at 95 °C with 5 min hold and two-segment cycles (95 °C with 15 s hold and 60 °C with 1 min hold) followed by a terminal step 72 °C.

### Genotyping and RT-PCR

Tail clippings from the tip of approximately 3 mm were collected from mice and used for DNA extraction (Roche, High Pure PCR Template Preparation Kit; manufacture’s protocol). For genotyping of Gemin4 genetrap (G4GT) the following primers were used. LTR-Forward: AAATGGCGTTACTTAAG-CTAGCTTGC, G4GT-Forward: GGAGCGAATATAGCCTTGATTCTCTGGAAATG, G4GT-Reverse: CTTCCCAGGACGGCCTCCTAGTCTTACCCTCTA.

The genotyping primers for the murine Smn Neo cassette are as follows. neostop-F: TCGCCTTCTTGACGAGTTCTTCTG, Smn-Forward: AGGATCTCTGTGTTCGTGCGTGGTG, Smn-Reverse: CCTTAAAGGAAGCCACAGCTTTATC. Dr. Cathleen Lutz of the Jackson Laboratory graciously supplied primer sequences for the *SMN2* transgene. PCR amplification was performed using standard protocols. For RT-PCR, total RNA was isolated from mouse tissues using Trizol reagent (Invitrogen; manufacture’s protocol). For RT-PCR analysis, the following primers were used: Gemin4 exon1-Forward – CAGACTACAGCACGGAAGCGGAG, Gemin4 exon2-Reverse – CTAAGCAGTTGGTGGTGCAGGATG.

### Western blotting

Protein lysates were prepared by flash freezing mouse tissues in liquid nitrogen then crushing the tissue into a fine powder. This powder was then transferred into RIPA-buffer with protease inhibitor (Thermo Scientific), homogenated with a 27G½ syringe and centrifuged at max speed at 4 °C for 10 min. Supernatants were quantified using a standard Bradford assay protocol. Equal amounts of samples (60 *μ*g) were subjected to 4 - 12 % MOPs/NuPage gradient gel system (Invitrogen) and transferred onto a nitrocellulose membrane (Schleicher & Schuell) following standard protocols. Lanes were transferred to nitrocellulose membranes and incubated with appropriate antibodies. Gemin4 immunoreactive bands were detected using the polyclonal Gemin4 Ab and horseradish peroxidase-conjugated goat antibodies to rabbit IgG (Gemin4 Ab 1:200 Santa Cruz Biotechnologies), monoclonal anti α tubulin Ab 1:5000 (Sigma-Aldrich) (secondary mouse Ab 1:10000, secondary rabbit Ab 1:5000, both from Thermo Scientific). Detection of protein bands was achieved using standard chemiluminescence substrates (SuperSignal West Femto, Thermo Scientific).

## Acknowledgments

We thank K.B. Shpargel for valuable help during early stages of the project and A.H. Natalizio for assistance with reformatting the figures. This work was supported by grants to A.G.M. from the NIH (R01-GM118636) and the Muscular Dystrophy Association (RG-4070). M.P.W. was supported in part by an NIH predoctoral traineeship (T32-GM08613).

---

**Figure S1.**
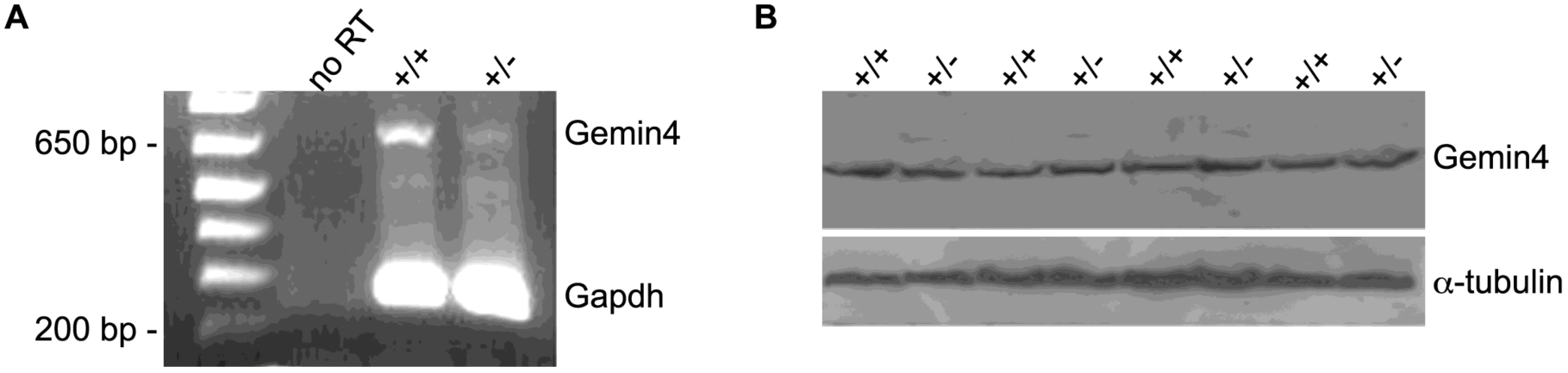
Expression of Gemin4 mRNA and protein. (A) RNA was isolated from livers of P4 *Gemin4^+/-^* and *Gemin4^+/+^* mice and semi-quantitative RT-PCR analysis was performed. Pooled *Gemin4^+/-^* and *Gemin4^+/+^* RNA for no RT control. (B) Western analysis of liver lysates was performed to determine protein levels of Gemin4 in wild-type and heterozygous animals; α-tubulin was used as a loading control.

